# MicroRNA exocytosis by large dense-core vesicle fusion

**DOI:** 10.1101/093278

**Authors:** Alican Gümürdü, Ramazan Yildiz, Erden Eren, Gökhan Karakülah, Turgay Ünver, Şermin Genç, Yongsoo Park

## Abstract

Neurotransmitters and peptide hormones are secreted into outside the cell by a vesicle fusion process. Although non-coding RNA (ncRNA) that include microRNA (miRNA) regulates gene expression inside the cell where they are transcribed, extracellular miRNA has been recently discovered outside the cells, proposing that miRNA might be released to participate in cell-to-cell communication. Despite its importance of extracellular miRNA, the molecular mechanisms by which miRNA can be stored in vesicles and released by vesicle fusion remain enigmatic. Using next-generation sequencing, vesicle purification techniques, and synthetic neurotransmission, we observe that large dense-core vesicles (LDCVs) contain a variety of miRNAs including miR-375. Furthermore, miRNA exocytosis is mediated by the SNARE complex and accelerated by Ca^2+^. Our results suggest that miRNA can be a novel neuromodulator that can be stored in vesicles and released by vesicle fusion together with classical neurotransmitters.

**One Sentence Summary:** Using next-generation sequencing (NGS) for microRNA (miRNA) and synthetic neurotransmission, we observed that large dense-core vesicles (LDCVs) contain a variety of miRNA together with classical neurotransmitters, and that miRNA can be released by vesicle fusion mediated by SNARE.

---

Non-coding RNA (ncRNA) is transcribed from the genome, but not translated into a protein. The protein-coding genes occupy ~1.2% of the euchromatic genome^1^, whereas the non-coding genes cover ~60% of the genome^2^. Approximately 98% of RNA transcripts are non-coding^2^. The main functions of ncRNAs are linked to translation, RNA splicing, and gene regulation^3^. MicroRNAs (miRNAs) are ncRNAs with 20~24 nucleotides in length; they are involved in a variety of cellular functions^4^, including gene expression and cellular communication^5^. miRNAs associate with the complementary sequences in target mRNAs^6^, and thereby repress mRNA translation.

Although miRNAs are present inside the cell where miRNAs are transcribed, miRNAs can be detected outside the cells, called extracellular miRNA^7, 8^. They participate in cell-to-cell communication^9^. Extracellular miRNAs have been hypothesized to be incorporated in secretory vesicles such as exosomes, microvesicles, and apoptotic bodies^10^, although the copy number is < 1^11^. Vesicle-free extracellular miRNAs are also detected in complexes with high-density lipoproteins (HDL)^12^ or Argonaute2^13^. More surprisingly, miRNAs seem to be released by active exocytosis in neurons^14–16^ and regulate pain signaling^17^. miRNAs including miR-29a and miR-125a can be secreted from synapses in a Ca^2+^-dependent manner^14^. Despite the increasing interest in extracellular miRNAs outside the neurons, little is known about the molecular mechanisms by which miRNAs can be released.

Neurons have two different types of vesicles that release neurotransmitters: synaptic vesicles and large dense-core vesicles (LDCV)^18, 19^. LDCVs are responsible for exocytosis of amines, neuropeptides, and hormones that mainly stimulate G-protein-coupled receptors (GPCRs)^18–20^. LDCVs, also called chromaffin granules, in chromaffin cells have been used as the model system for LDCVs^21^. LDCVs and synaptic vesicles share the conserved fusion machinery; i.e., soluble *N*-ethylmaleimide-sensitive factor attachment protein receptor (SNARE) proteins and Ca^2+^ sensor synaptotagmin-1 that mediates Ca^2+^-dependent exocytosis ^22^.

To study miRNA exocytosis by vesicle fusion, we focus on LDCVs because of their large size, high yield, and high purity. We purified LDCVs (95% purity) from bovine chromaffin cells, then used next-generation sequencing (NGS) to sequence miRNAs stored in LDCVs. Synthetic neurotransmission, the *in-vitro* reconstitution of vesicle fusion, supported the hypothesis that miRNA exocytosis is mediated by SNARE proteins and accelerated by Ca^2+^. These results suggest that miRNAs may be neuromodulators stored in vesicles and released by vesicle fusion along with classical neurotransmitters.

## Results

We have established the LDCV purification technique based on continuous sucrose gradients using bovine adrenal medulla as described before^23, 24^. Vesicle proteins are highly enriched, whereas other contaminants are mostly removed^23^. Mitochondria, late, lysosomes, early endosomes, endoplasmic reticulum, proteasomes, and peroxisomes are not detected in LDCV samples^23^. This purification process yields primarily mature LDCVs with 100 ~ 200 nm in diameter^23^; i.e., immature LDCVs (a marker, VAMP-4) are mostly removed.

We confirmed the enrichment of LDCVs by western blotting showing that LDCVs are highly enriched in vesicle membrane proteins (synaptotagmin-1 and synaptobrevin-2/VAMP-2), whereas exosomes and multivesicular bodies (a marker, LAMP-1) are removed during purification. (**Supplementary Fig. 1**). Proteomic profiling of exosomes^25^ and western blotting data^26^ reveal that exosomes are enriched with LAMP-1. Thus, our data support that miRNA-containing organelles are excluded.

To quantify the LDCV purity, we performed an overlay assay (**Fig. 1a-d**) in which LDCVs were immobilized on a poly-L-lysine-coated glass slides, then immunostained with an antibody against synaptobrevin-2/VAMP-2 (a marker for LDCVs, green) and DiI (a lipophilic membrane dye, red). Overlaps (yellow) in an overlay assay indicate LDCVs, whereas contaminants stain red; note that mitochondria could be minorly included in LDCV samples^23^. An overlay assay shows 95% purity of LDCVs (119 LDCVs from 125 samples, Fig. 1d). Size distribution of purified LDCVs with average of 135 nm determined by dynamic light scattering (Fig. 1e) correlate with electron microscopy data previously reported^23^.

**Figure 1.**
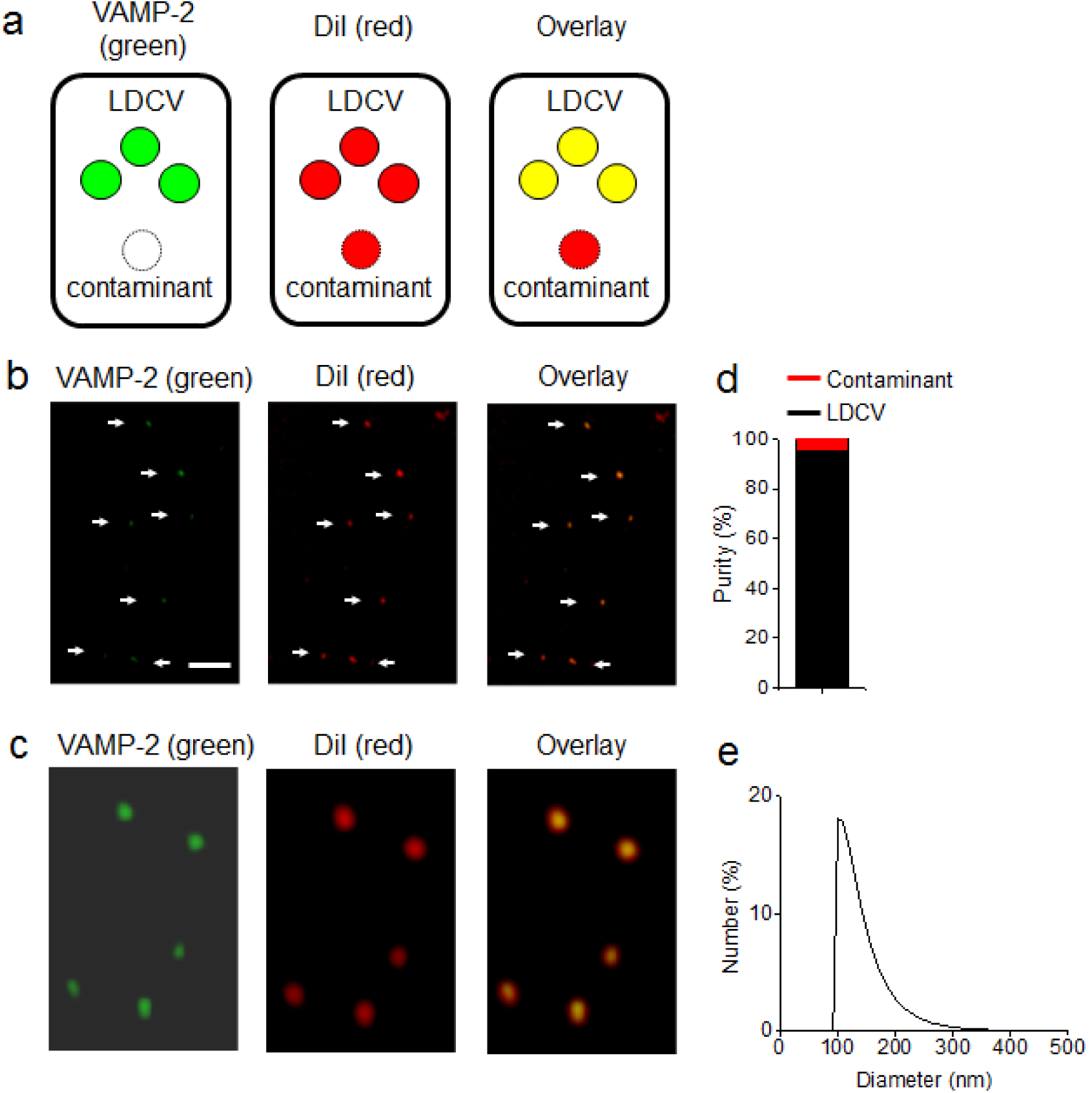
High purity of LDCVs. (**a**) Schematic of the overlay assay for the LDCV purity. (**b-c**) LDCVs were incubated simultaneously with VAMP-2 antibody (green, left) and a membrane dye, DiI (red, middle). Overlay (yellow, right) indicates LDCVs (arrows). Scale bar, 2 μm. (**c**) High-magnification images of LDCVs stained with VAMP-2 antibody and DiI. (**d**) LDCV purity was presented as a percentage of LDCVs (yellow from panel **b-c**, *n* = 119) among total vesicles (red in overlay from panel **b-c**, *n* = 125). Total vesicles were collected from three independent purifications. The purity of LDCVs is 95%. (**e**) Size distribution of LDCVs analyzed using dynamic light scattering. Number distribution is presented as a percentage.

Next, total RNAs were extracted from purified LDCVs by using miRNeasy Mini Kit (Qiagen). The RNAs stored in LDCVs were < 100 nucleotides (nt) long (Fig. 2a); rRNA was not detected, although it constitutes ~80% of the total RNA in a cell, suggesting that LDCVs are specifically enriched with small RNAs. The small RNA libraries (**Supplementary Fig. 2**) were read using small RNA-sequencing (RNA-seq) technique with Illumina Hiseq 2500 platform. Surprisingly, the RNA library provides evidence for miRNA fraction at the peak with 141 bp; miRNAs of 20 ~ 24 nt are conjugated to ~120 bp of adapter (**Supplementary Fig. 2**). miRNA fraction constitutes ~61% of RNAs extracted from LDCVs; i.e., miRNAs are selectively accumulated in LDCVs (Fig. 2b). Size selection of miRNA fraction was applied for miRNA sequencing and we found a variety of miRNAs including 459 of known miRNAs and 315 of novel miRNAs predicted by miRDeep2 analysis tool (Fig. 2c) (**Supplementary File 1,2**). However, most of the known miRNAs were expressed at low level less than 10 (on a log2 scale) of library size normalized Count Per Million (CPM) (Fig. 2d). Therefore, we used a criterion of 1,000 CPM (9.96 on a log2 scale) to categorize miRNAs as high (69 miRNAs, 16.2%) and low (356 miRNAs, 83.8%) level miRNA (Fig. 2e,f). Intriguingly, the 69 high level miRNAs (Fig. 2e) comprise 94.8% of total mapped read counts (Fig. 2f); i.e., only a few miRNAs are highly expressed in LDCVs. miR-375 was the most abundant miRNA in LDCVs (28.9% of total mapped read counts, Fig.2g). The most abundant miRNAs (top 5%; 21 miRNAs among 425 total miRNAs) stored in LDCVs account for 80% of CPM (Table 1). We confirmed the reproducibility of RNA-seq data by using technical and biological replicate which are highly correlated (r=0.99, p<0.05) (**Supplementary Fig. 3**). These data provide evidence that miRNAs including miR-375 are highly enriched in LDCVs.

**Figure 2.**
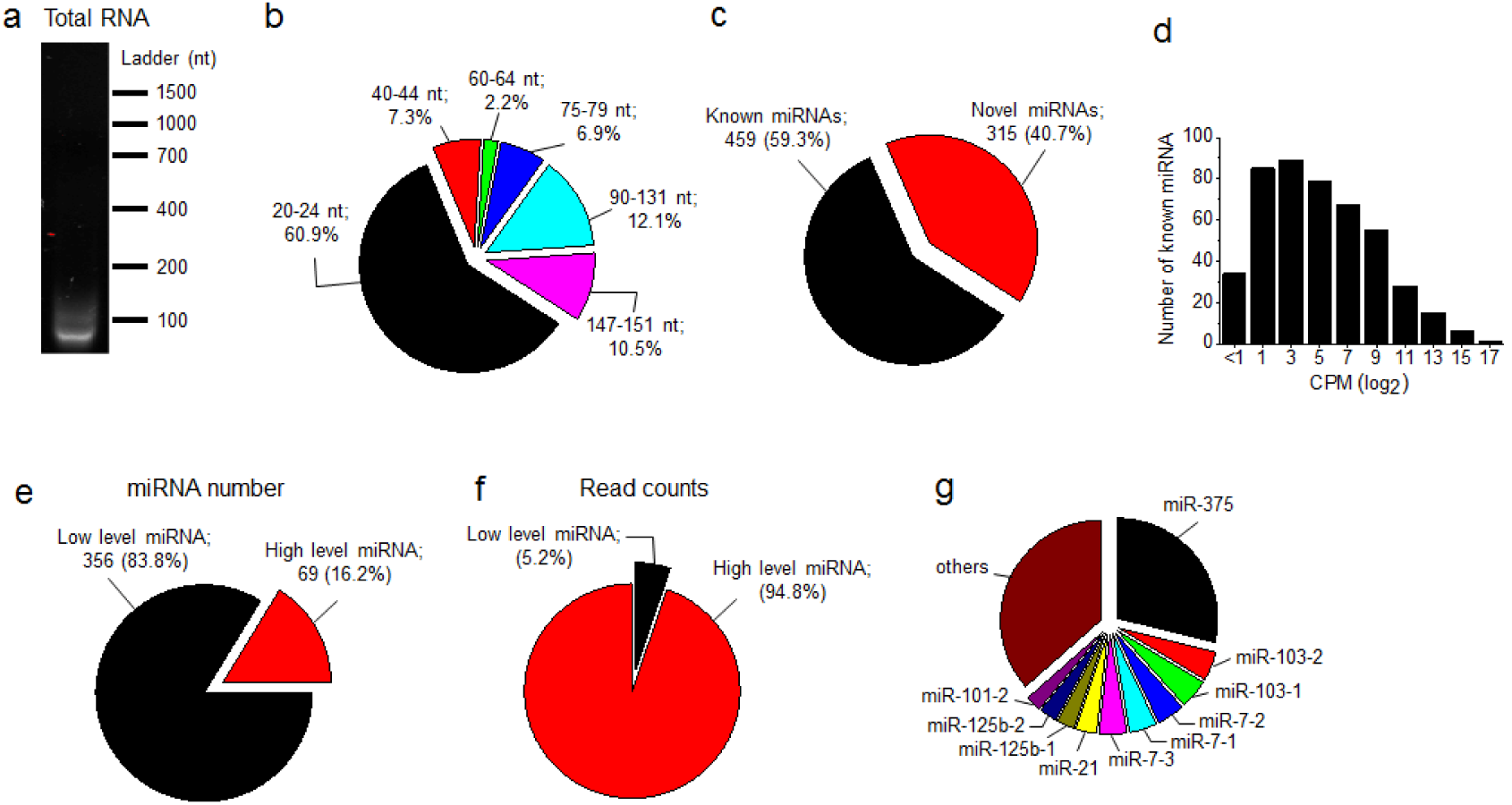
Next generation sequencing for miRNAs. (**a**) Total RNA extracted from LDCVs was analyzed by electrophoresis on a 2% agarose gel. (**b**) Size distribution of the small RNAs. Values were calculated as a percentage based on results of electropherogram trace of total RNA libraries in the **Supplementary Fig. 2**. (**c**) Known miRNA and novel miRNA based on the number of miRNAs from RNA-seq. (**d**) Distribution of the number of miRNAs based on read counts; count per million (CPM) on a log_2_ scale. Number of known miRNAs in each CPM (log_2_) range is presented. (**e,f**) Classification of the number of miRNAs as high level (69 miRNAs) and low level (356 miRNAs) based on read counts; 1,000 CPM was applied for this classification after cut-off of < 1 log_2_ CPM (**e**). (**f**) High-level and low level-miRNAs were presented as a percentage of read counts, CPM. (**g**) Distribution of the most abundant miRNAs based on CPM.

**Table 1.** Top 5% of the most abundantly expressed known miRNAs stored in LDCVs

| miRNA ID | Mature miRNA accession ID | miRBase precursor id | Count per million (CPM) | % total CPM | Chromosome | Start locus | Stop locus | Strand |
| --- | --- | --- | --- | --- | --- | --- | --- | --- |
| bta-miR-375 | MIMAT0009303 | MI0009817 | 288587.23 | 28.9 | chr2 | 107667521 | 107667584 | - |
| bta-miR-103 | MIMAT0003521 | MI0005456 | 47276.18 | 4.7 | chr13 | 51742422 | 51742497 | - |
| bta-miR-103 | MIMAT0003521 | MI0004736 | 47272.63 | 4.7 | chr20 | 189857 | 189928 | - |
| bta-miR-7 | MIMAT0003843 | MI0010462 | 44812.40 | 4.5 | chr8 | 78575531 | 78575639 | - |
| bta-miR-7 | MIMAT0003843 | MI0010471 | 44751.70 | 4.5 | chr21 | 19989064 | 19989161 | + |
| bta-miR-7 | MIMAT0003843 | MI0005056 | 44739.55 | 4.5 | chr7 | 20584157 | 20584238 | - |
| bta-miR-21 | MIMAT0003528 | MI0004742 | 34130.20 | 3.4 | chr19 | 11033072 | 11033143 | + |
| bta-miR-125b | MIMAT0003539 | MI0005457 | 28765.84 | 2.9 | chr1 | 19881347 | 19881431 | - |
| bta-miR-125b | MIMAT0003539 | MI0004753 | 28774.84 | 2.9 | chr15 | 33298815 | 33298902 | - |
| bta-miR-101 | MIMAT0003520 | MI0004735 | 25492.58 | 2.5 | chr8 | 39940832 | 39940910 | - |
| bta-miR-101 | MIMAT0003520 | MI0009721 | 25420.41 | 2.5 | chr3 | 80666417 | 80666499 | + |
| bta-miR-27b | MIMAT0003546 | MI0004760 | 18647.03 | 1.9 | chr8 | 83009823 | 83009919 | + |
| bta-miR-26a | MIMAT0003516 | MI0004731 | 17461.18 | 1.7 | chr5 | 55977923 | 55978006 | + |
| bta-miR-26a | MIMAT0003516 | MI0009784 | 17457.90 | 1.7 | chr22 | 11457900 | 11457989 | + |
| bta-miR-30d | MIMAT0003533 | MI0004747 | 14900.95 | 1.5 | chr14 | 8080292 | 8080361 | + |
| bta-miR-10b | MIMAT0003839 | MI0005052 | 13952.60 | 1.4 | chr2 | 20797606 | 20797704 | - |
| bta-miR-504 | MIMAT0009340 | MI0009855 | 12472.64 | 1.2 | chrX | 22083689 | 22083771 | - |
| bta-miR-127 | MIMAT0003787 | MI0005008 | 11830.13 | 1.2 | chr21 | 67429744 | 67429838 | + |
| bta-miR-379 | MIMAT0009306 | MI0009820 | 11663.16 | 1.2 | chr21 | 67561869 | 67561954 | + |
| bta-miR-411a | MIMAT0009312 | MI0009826 | 10604.45 | 1.1 | chr21 | 67563098 | 67563179 | + |
| bta-miR-143 | MIMAT0009233 | MI0009743 | 10375.68 | 1.0 | chr7 | 62809304 | 62809404 | + |

Can miRNAs stored in LDCVs be released to extracellular fluid? We attempted to confirm miRNA exocytosis by using the reconstitution system of vesicle fusion called synthetic neurotransmission. We used purified LDCVs for synthetic neurotransmission and prepared plasma membrane-mimicking liposomes that contain the stabilized Q-SNARE complex (**Online Methods**). Liposomes also incorporated GelGreen dye, a water-soluble and membrane-impermeable green fluorescent nucleic acid dye. As a content-mixing assay, miRNAs stored in LDCVs bind to GelGreen after LDCV fusion, thereby increasing fluorescence of GelGreen (Fig. 3a). Indeed, we observed miRNA exocytosis mediated by SNARE assembly, because soluble VAMP-2 or no SNARE proteins in plasma membrane-mimicking liposomes completely inhibited miRNA exocytosis (Fig. 3b). 100 μ M Ca^2+^ accelerated miRNA exocytosis that resulted from the activity of synaptotagmin-1, a Ca^2+^ sensor, for vesicle fusion (Fig. 3b, **Supplementary Fig. 4**). To further validate that miR-375 can be released from cells in response to stimulation, we used PC12 cell, the cell line derived from a pheochromocytoma of chromaffin cells. Indeed, stimulation of PC 12 cells with 50 mM KCl that evokes membrane depolarization and Ca^2+^ influx increased the level of miR-375 in the extracellular (Fig. 3c), supporting miR-375 exocytosis by vesicle fusion in the physiological conditions.

**Figure 3.**
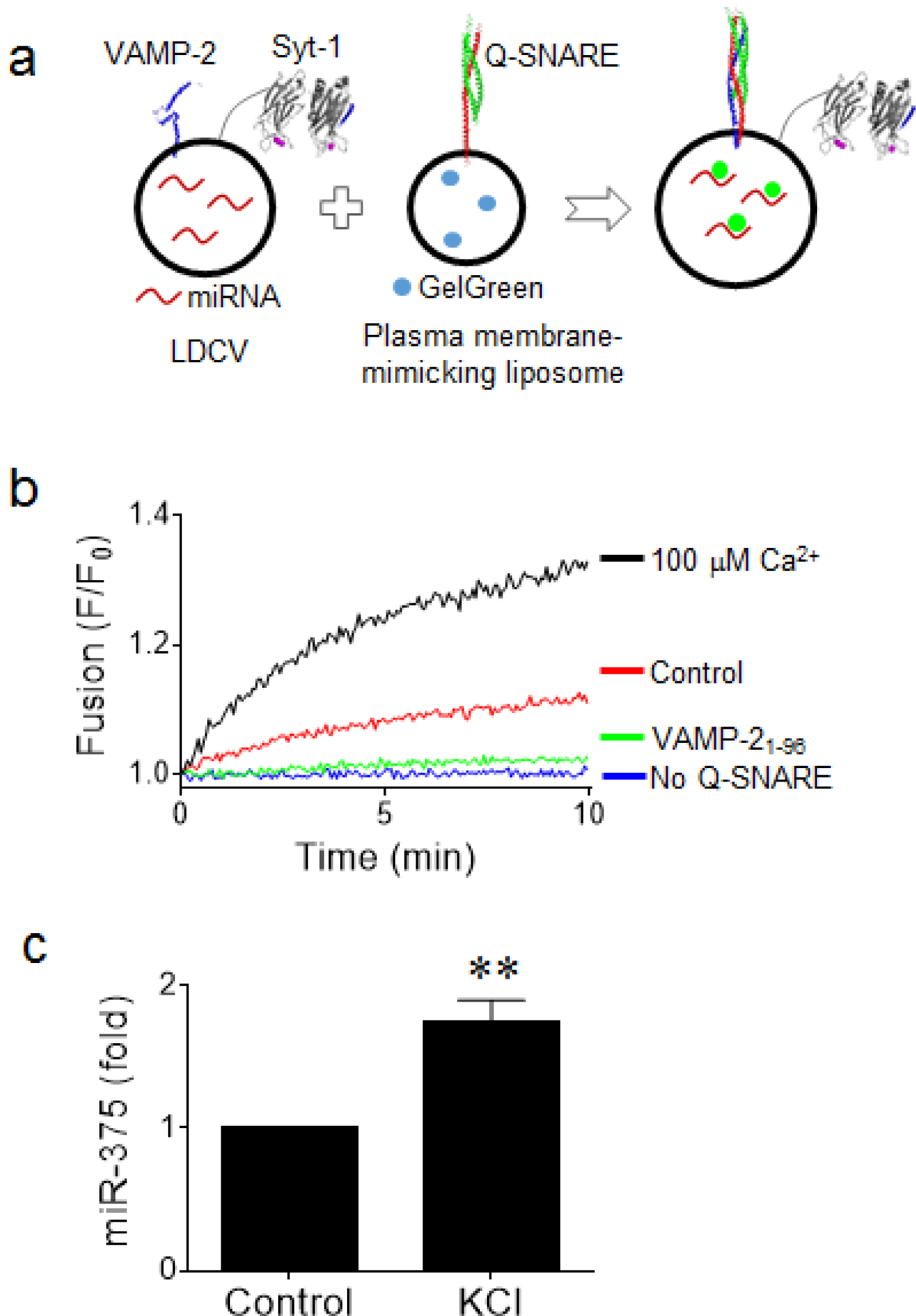
miRNA exocytosis by vesicle fusion. (**a**) Schematic for fusion assay. Fusion was monitored using a content-mixing assay in which GelGreen, a water-soluble and membrane-impermeable nucleic acid dye, was incorporated in liposomes. Interaction of miRNA stored in LDCVs with GelGreen increases its fluorescence. Plasma membrane-mimicking liposomes incorporated the stabilized Q-SNARE complex^45^. VAMP-2 fragment is omitted for clarity. (**b**) Preincubation of liposomes with a soluble fragment of VAMP-2 (VAMP-2_1-96_), inhibits SNARE-mediated fusion due to competitive inhibition^24, 46^. Control, basal fusion of LDCVs with liposomes that contain Q-SNARE. No Q-SNARE, no SNAREs incorporated in liposomes. Fluorescence intensity is normalized to the initial value (F_0_). (**c**) miR-375 exocytosis from PC12 cells. PC12 cells were stimulated with 50 mM KCl for 5 min. miR-375 released by PC 12 cells in the absence or presence of KCl was analyzed using qRT-PCR. Quantitative analysis was presented as fold changes in expression relative to a control (no KCl). Data are mean ± SD (*n* = 4 biological and technical replicates). **, *p* = 0.0026 (twotailed unpaired Student’s t test).

Next, we quantified the copy number of miR-375 per LDCV (Fig. 4a,b). The number of LDCV particles was measured using Nanoparticle Tracking Analysis (NTA). To determine the copy number of miR-375, which is the most abundant miRNA in LDCVs, we used quantitative real-time PCR (qRT-PCR) to quantify miR-375. A synthesized *Caenorhabditis elegans* microRNA (cel-miR-39) as the spike-in control was added to normalize the RNA extraction efficiency. Using the data derived from the above experiments, we analyzed the relative ratio of miR-375 to LDCVs for each sample. Each LDCV contained approximately 400 ~ 500 miR-375 (Fig. 4b), suggesting that LDCVs contain much higher number of miRNAs than other miRNA-containing organelles such as exosomes^11^.

**Figure 4.**
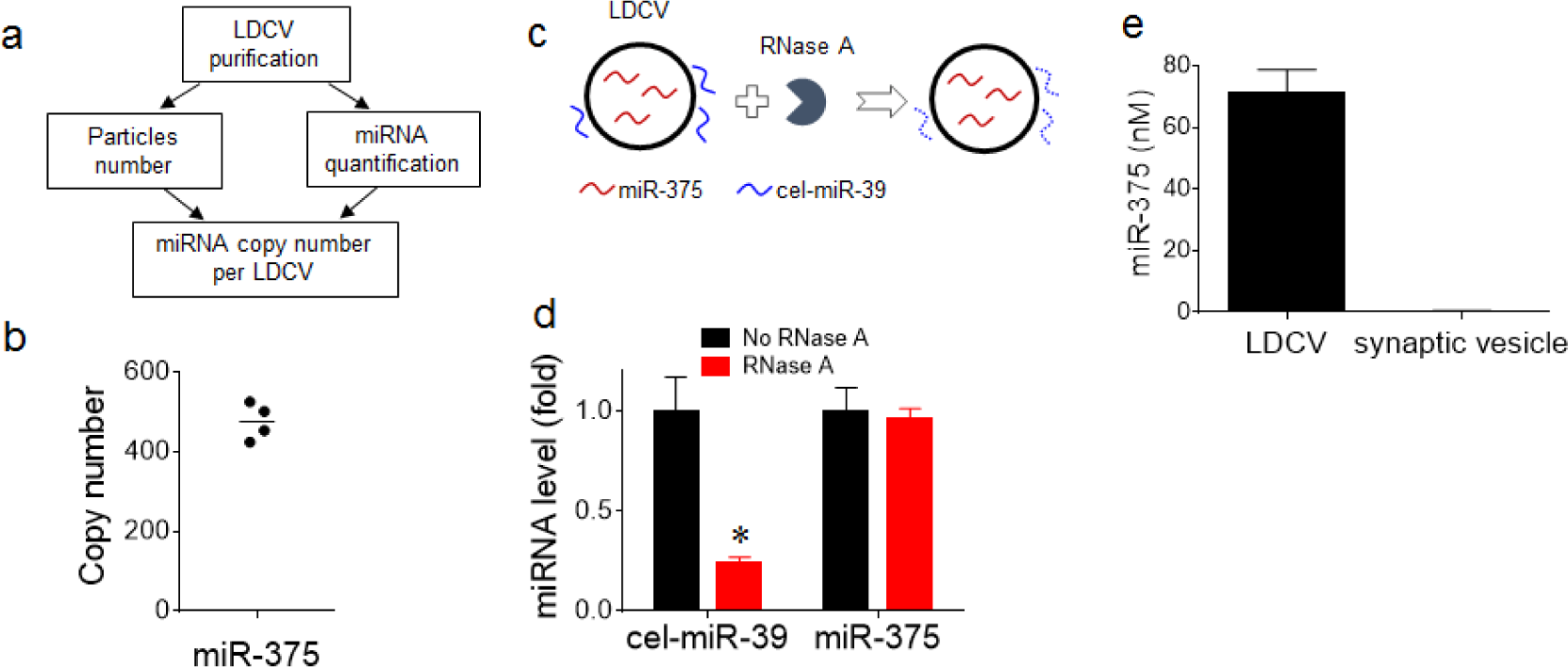
miR-375 stored in LDCVs. (**a,b**) Quantification of miRNA copy number per LDCV. (**a**) Workflow to determine miRNA copy number. Aliquots of LDCVs were counted using Nanoparticle Tracking Analysis (NTA). In parallel, the concentration of miRNA extracted from LDCVs was determined by qRT-PCR. A synthesized *Caenorhabditis elegans* microRNA (cel-miR-39) as the spike-in control was added to normalize the RNA extraction efficiency. (**b**) Copy number of miR-375 per LDCV. Line represents mean (*n* = 4 biological and technical replicates). (**c**) Schematic for confirming vesicle-incorporated miR-375. cel-miR-39 was mixed with LDCVs prior to RNase A treatment. (**d**) Relative levels of cel-miR-39 and miR-375 with or without RNase A were determined by qRT-PCR. Values represent fold changes in expression relative to a control (no RNase A). *, *p* = 0.014 (two-tailed unpaired Student’s t test). (**e**) miR-375 from purified synaptic vesicles from mice brains was analyzed by qRT-PCR. Data in **d** and **e** are mean ± SD (*n* = 3 technical replicates).

Can miR-375 be a contaminant from the cytosol, which is adsorbed to the outside of LDCVs during purification procedure? Next, we assessed the possibility that miR-375 could attach to vesicle membranes during the purification process as an artefact (Fig. 4c,d). To confirm that miR-375 is really stored inside LDCVs, we applied RNase A, because RNase A selectively degrades vesicle-free miRNA, but not vesicle-incorporated miRNA. Indeed, RNase A degraded cel-miR-39, which was mixed with LDCVs prior to RNase A, whereas miR-375 was intact (Fig. 4d), indicating that miR-375 is incorporated in LDCVs. Taken together, our data suggest that miR-375 is stored in LDCVs (~500 copies) and miRNA exocytosis is mediated by SNARE assembly and accelerated by Ca^2+^ stimulus.

Finally, we tested the specificity of miR-375 by investigating whether miR-375 is present in synaptic vesicles purified from mice brains (Fig. 4e). Surprisingly, we found that miR-375 is only stored in LDCVs, but not in synaptic vesicles, supporting the specificity of miR-375. Altogether, these data provide evidence that miR-375 is not an artefact or a byproduct caused by vesicle purification procedure, but miR-375 might be a novel neuromodulator stored in LDCVs and released by vesicle fusion in response to stimulation.

## Discussion

Our data strongly suggest that miRNAs are stored in vesicles and can be released along with classical neurotransmitters by vesicle fusion. LDCVs contain non-coding RNA (ncRNA) together with catecholamines. Exosomes also carry ncRNAs, but up to 90% of these are rRNAs and miRNAs are less than 5%^27^. Furthermore, abundant miRNAs are present at less than one copy per exosome^11^. In comparison with exosomes, the variety of miRNAs is highly enriched in LDCVs excluding other types of ncRNA such as rRNAs and tRNAs. These observation suggests that LDCVs may be specific cargos and carriers in neurons and neuroendocrine cells that deliver miRNAs to extracellular fluid.

The average diameter of LDCVs is 167.7 nm^23^, with inner radius of 80 nm, which corresponds to 2.14 x 10^-18^ liter. Therefore, 400 ~ 500 copies of miR-375 in a single LDCV is a concentration of 320 ~ 400 μM. LDCV in chromaffin cells contains ~2.5 x 10^6^ catecholamine molecules per vesicle^28^ and a variety of neuropeptides; e.g., ~430 neuropeptide Y (NPY) molecules^21, 29^. Therefore, the observed miRNA concentration is comparable to the number of other neuropeptides in LDCVs, proposing that miRNAs might be novel neuromodulators or hormones that are stored in LDCVs and released together with catecholamines and neuropeptides (Fig. 5). Therefore, we propose a new function of noncoding RNAs named ‘ribomone <u>(ribo</u>nucleotide + ho<u>rmone)</u>’ that are stored inside the vesicles and released by active vesicle fusion in response to stimulation, thereby regulating cell-to-cell communication including gene silencing and cellular signaling.

**Figure 5.**
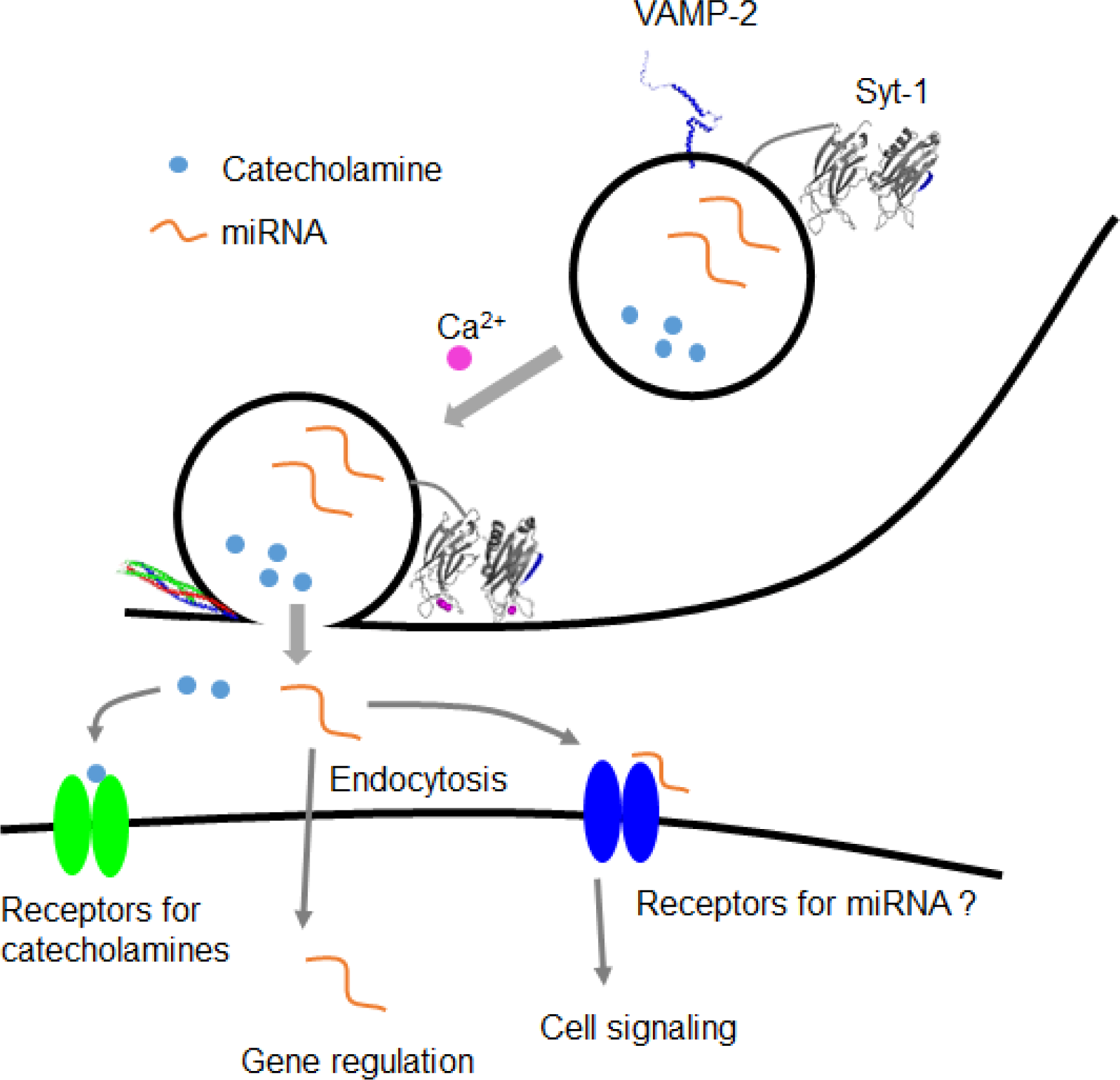
A schematic summarizing miRNA exocytosis by vesicle fusion. LDCVs contain catecholamines that include dopamine, adrenaline, and noradrenaline, but they also contain a variety of miRNAs including miR-375. miRNA exocytosis is mediated by the SNARE complex and accelerated by Ca^2+^ stimulus. Released extracellular miRNAs regulate cell-to-cell communication by controlling gene silencing in target cells after endocytosis and by stimulating receptors or ion channels, thereby leading to cellular signaling. Our data suggest the new concept that miRNAs are stored in vesicles together with classical neurotransmitters and are released by vesicle fusion, thus contributing to cell-to-cell communication as a novel neuromodulator.

Extracellular circulating miRNAs have been detected in serum, so they might be biomarkers of diseases ^10, 30, 31^. Most studies have proposed that extracellular miRNAs can be packaged into membrane-associated organelles such as exosomes, microvesicles, and extracellular vesicles^10, 32^, but the mechanisms and origins of vesicle-free extracellular miRNAs remain unclear. To our knowledge, our data are the first to identify miRNA exocytosis by LDCV fusion. LDCVs in chromaffin cells, which are neuroendocrine and modified sympathetic ganglion cells, mainly contain catecholamines^33^. Upon activation of the sympathetic nervous system, catecholamines released by LDCV fusion spread through the bloodstream, thereby mediating stress and the ‘fight-or-flight’ physiological response^33^. Therefore, fusion of LDCVs could result in a high level of vesicle-free miRNAs together with catecholamines in blood.

miR-375 is the most abundant miRNA in LDCVs (~30% of total miRNA read counts); it is highly expressed in the pancreatic islets^34^ and motor neurons^35^, where it regulates neuronal development^36^. miR-375 is also detected in plasma and serum as a circulating miRNA, and may be a biomarker for diabetes^37^, hepatocellular carcinoma^38^, and Alzheimer’s disease^39^. Despite intense study of miR-375, the physiological roles and origin of circulating extracellular miR-375 are still not understood. Our data support the hypothesis that LDCVs in chromaffin cell may be one source of circulating miR-375, but its function remains to be elucidated in both normal physiological and pathological conditions.

Besides miR-375, LDCVs are enriched with miR-7, miR-103, and miR-125b, which are highly expressed in the brains^40, 41^. miR-7 regulates neuronal development^42^ and miR-125b promotes neuronal differentiation^43^. miR-29a and miR-125a are secreted from synaptosomes in response to depolarization^14^, and both miR-29a and miR-125a were among the 10% of the most abundant miRNAs in LDCVs (**Supplementary File 3**). Another miRNA, let-7b, is released from dorsal root ganglion (DRG) neurons by depolarization and extracellular let-7b activates toll-like receptor-7 (TLR7)/TRPA1 ion channel to mediate pain signaling^17^. let-7b was among the 15% most-abundant miRNAs in LDCVs (**Supplementary File 3**).

We have essential open questions. First, what are miRNAs doing after release and what is the physiological significance? Extracellular circulating miRNAs are aberrantly detected in blood plasma and serum^7, 8^, probably regulating cell-to-cell communication^9^. Extracellular miRNAs could be transported by endocytosis into the target cells, where they regulate gene expression, but the identity of the target cells and the mechanisms of endocytosis are not fully understood. Secondly, miRNA clearly regulates gene expression inside the cell, but we hypothesize that extracellular miRNAs may directly regulate receptor activity as an agonist or antagonist, similarly to aptamers, which are oligonucleotides that bind to a specific target, in a manner that differs from the way in which it regulates gene expression (Fig. 5). For example, extracellular let-7b miRNA activates TLR7 to mediate pain signaling^17^. So far, little is known about the signaling mechanism by which extracellular miRNAs can regulate receptors and ion channels. Thirdly, the mechanism by which miRNAs are transported into vesicles is not known. Considering that some miRNAs (e.g., miR-375) are selectively and specifically accumulated in LDCVs, LDCVs could have a carrier or transporter for miRNA uptake. LDCVs express nucleotide transporters^44^, but the activity of RNA uptake has never been studied. Fourthly, ribonucleases in the extracellular could degrade vesicle-free extracellular miRNAs. High-density lipoproteins (HDL)^12^ or Argonaute2^13^ might protect extracellular miRNAs from cleavage by ribonucleases. We have the mass spectrometry data showing that RNA-binding proteins such as hnRNPs and apolipoprotein A-I are detected in LDCVs whereas Argonaute2 is not (unpublished data). Further work will be required to identify miRNA-binding proteins that control the miRNA stability and miRNA function in LDCVs.

## Acknowledgments

We thank Dr. Reinhard Jahn for all constructs and samples. We are deeply indebted to Dr. Kyong-Tai Kim for technical support. We thank the Sequencing Service GeneCore Sequencing Facility (EMBL, http://www.genecore.embl.de) for RNA-seq analysis. This work was supported by EMBO Installation Grant (IG Project Number 3265 to Y.P) and 2236 COFUNDED BRAIN CIRCULATION SCHEME (Project Number 116C022 to Y.P).

## Author Contributions

A.G., R.Y, and E.E. performed RNA extraction from LDCVs, synaptic vesicles, and PC12 cells. A.G., R.Y., and E.E. performed RNA preparation for NGS. A.G., R.Y., and E.E. carried out absolute miRNA quantification for counting copy number and confirming vesicle-incorporated miR-375. A.G., R.Y., and Y.P. contributed fusion assay. A.G. carried out dynamic light scattering and bioanalyzer. G.K. and T. U. performed computational analysis of RNA-sequencing data. Y.P. carried out LDCV purification and overlay assay. Y.P. designed, collected, and analyzed data. S.G. and Y.P. wrote the manuscript and all authors read and provided their comments on the draft. Y.P. supervised the project.

## Author information

Our NGS data (RNA-Seq) have been deposited with Gene Expression Omnibus (GEO) under accession codes GSE84834, GSM2252051 (miRNAs_LDCV_technical replicate_1), GSM2252052 (miRNAs_LDCV_technical replicate_2), and GSM2252053 (miRNAs_LDCV_biological replicate_1). The following link has been created to allow review of record GSE84834 while it remains in private status:http://www.ncbi.nlm.nih.gov/geo/query/acc.cgi?token=wzwhakcynvalfcv&acc=GSE84834

The authors declare no competing financial interests. Correspondence and requests for materials should be addressed to Y.P.

## Online Methods

### Materials

Poly-L-lysine was from sigma (Cat: P8920) and DiIC18 was purchased from Invitrogen (Cat: D-12730, Disulfonic Acid). Monoclonal mouse antibody against synaptobrevin-2/VAMP-2 (clone, 69.1) and synaptotagmin-1 (clone, 41.1) were obtained from Synaptic Systems (Gottingen, Germany). LAMP-1 (Abcam, cat. no. ab24170) and Goat Anti-Mouse IgG (Cy2) (ab6944) was obtained from Abcam (Cambridge, UK). DC (detergent compatible) protein assay was obtained from Bio-Rad (DC Protein Assay, Cat: 5000112). GelGreen was purchased from Biotium (Hayward, CA, Cat: 41005). Nucleospin RNA plus for total RNA extraction was obtained from MACHEREY-NAGEL (Duren, Germany, Cat: 740984).

### Purification of large dense-core vesicles (LDCVs) and synaptic vesicle

LDCVs were purified using a previously-reported method^47^ with several modifications^23^. Fresh bovine adrenal glands were obtained from a local slaughterhouse. The cortex and fat were removed, then the medullae were minced using scissors, then homogenized in a cooled glass-teflon homogeniser. Crude LDCV fraction was resuspended in 300 mM sucrose buffer and loaded on top of a continuous sucrose gradient (from 300 mM to 2.0 M) to remove other contaminants including mitochondria. LDCVs were collected from the pellet after centrifugation at 110,000 g for 60 min in a Beckman SW 41 Ti rotor and resuspended with the buffer (120 mM K-glutamate, 20 mM K-acetate, 20 mM HEPES.KOH, pH 7.4). Protein concentration of LDCVs were determined using a Lowry assay (Bio-Rad DC Protein Assay, Cat: 5000112). Size distribution of purified LDCVs was determined using dynamic light scattering (NanoPlus DLS, Particulate Systems).

Synaptic vesicles from mice brain were purified as described in detail elsewhere^48^. Briefly, mice brains were homogenized in homogenization buffer supplemented with protease inhibitors, using a glass-Teflon homogenizer. The homogenate was centrifuged for 10 min at 1,000 g and the resulting supernatant was further centrifuged for 15 min at 17,000 g. The synaptosome pellet was lysed by adding ice-cold water, followed by the centrifuge for 25 min at 48,000 g. The resulting supernatant was overlaid onto a 0.7 M sucrose cushion and centrifuged for 1 h at 133,000 g. The pellet was resuspended in the buffer (120 mM K-glutamate, 20 mM K-acetate, 20 mM HEPES.KOH, pH 7.4).

### PC12 cell culture

The cells were maintained in RPMI-1640 Medium (Biochrom GmbH, Germany) containing 10 % horse serum, 5% fetal bovine serum (Gibco), and 1% antibiotics (Biochrom GmbH, Germany). Cells were cultured in T-25 flask (Sarstedt, Germany) and used for experiments at 60~70% of confluency. After washing twice with PBS supplemented with 1 mM Mg_2_Cl and 2 mM Ca_2_Cl, PC 12 cells were stimulated with 50 mM KCl for 5 min. Extracellular buffer was taken and subjected to qRT-PCR after centrifuging at 1,200 g for 5 min to remove cell debris followed by RNA isolation.

### Protein purification

All SNARE constructs were based on *Rattus norvegicus* sequences. The stabilized Q-SNARE acceptor complex consisting of syntaxin-1A (aa 183-288), SNAP-25A (aa 1-206, no cysteine), and a C-terminal VAMP-2 fragment (aa 49-96) was purified as described earlier^23, 45^. The stabilized Q-SNARE complex was expressed using cotransformation^24^ and purified by ion-exchange chromatography on Mono Q column (GE Healthcare, Piscataway, NJ) in the presence of 50 mM n-octyl-β-D-glucoside.

Protein structures were visualized using the program PyMOL (PDB ID: 1BYN for C2A, 1K5W for C2B, 3IPD for the SNARE complex, 2KOG for VAMP-2).

### Preparation of plasma membrane-mimicking liposomes

Lipid composition (molar percentages) of liposomes that contain the Q-SNARE complex consists of 45% PC (L-a-phosphatidylcholine), 15% PE (L-α-phosphatidylethanolamine), 10% PS (L-α-phosphatidylserine), 25% Chol (cholesterol), 4% PI (L-α-phosphatidylinositol), and 1% PIP_2_. Lipid mixture dissolved in chloroform/methanol (2:1, v/v) was dried under a gentle stream of nitrogen in the hood for 5 min, thereby giving rise to a lipid film. The lipid film was resuspended with 25 μL buffer containing 150 mM KCl, 20 mM HEPES/KOH pH 7.4, and 5% sodium cholate. In parallel, the Q-SNARE complex was resuspended in 75 μL buffer containing 150 mM KCl, 20 mM HEPES/KOH pH 7.4, and 1.5% sodium cholate. For a content-mixing assay, GelGreen, a water-soluble and membrane-impermeable nucleic acid fluorescent dye, was included in buffer in 1:100 dilution. The protein and lipid samples were mixed in protein-to-lipid molar ratio of 1:500 (100 μL in total), then a size exclusion column was used to remove detergent (Sephadex G50 in 150 mM KCl and 20 mM HEPES, pH 7.4).

Plasma membrane-mimicking liposomes that contain the Q-SNARE complex were collected as eluted; note that plasma membrane-mimicking liposomes incorporate GelGreen dye.

### LDCV fusion assay

LDCV fusion *in vitro* was monitored using a content-mixing assay in buffer containing 120 mM K-glutamate, 20 mM K-acetate, 20 mM HEPES-KOH (pH 7.4), 1 mM MgCl_2_, and 3 mM 2Na-ATP. LDCVs and plasma membrane-mimicking liposomes were incubated and interaction of miRNAs with GelGreen induced by vesicle fusion increase fluorescence of GelGreen (Ex: 495 nm/Em: 520 nm). 100 μM free Ca^2+^ concentrations in the presence of ATP and Mg^2+^ were calibrated using the Maxchelator simulation program (http://maxchelator.stanford.edu).

### Overlay assay for the LDCV purity

LDCVs (5 μg) were incubated on glasses (18 mm in diameter) coated with 0.1% poly-L-ly sine for 30 min at room temperature (RT) in 150 μL buffer containing 150 mM KCl and 20 mM HEPES, pH 7.4 with KOH. After fixing LDCVs with 4% PFA (in PBS) at RT for 15 min, then PFA was removed and LDCVs were washed three times with PBS. Monoclonal mouse antibody against synaptobrevin-2/VAMP-2 (Synaptic Systems; clone, 69.1) was incubated for 30 min at RT and then Cy2-linked goat anti-mouse IgG (Ex/Em: 489/505 nm) was incubated for 30 min at RT. LDCVs were simultaneously incubated with 1 μM DiI (Ex/Em: 549/565 nm) with VAMP-2 antibody. DiI stains membranes of LDCVs and other contaminant organelles. The two channels (green and red) constitute an overlay images through direct observation in a microscope. The LDCV purity was presented as the percentage of the number of LDCVs from the organelles counted (n = 125).

### RNA-seq data

RNA sequencing (RNA-seq) was performed by GeneCore Sequencing Facility (EMBL, http://www.genecore.embl.de) using Illumina Hiseq 2500. Briefly, total RNA (~1 μg) was extracted from ~300 μg of purified LDCVs by using miRNeasy Mini Kit (Qiagen, Cat: 217004). Then, the small RNA libraries have been prepared starting with 500 ng of RNA using the NEBNext^®^ Multiplex Small RNA Library Prep Set for Illumina^®^ (Set 1) (NEB, Cat: E7300S/L). Size selection of the fraction around 140 bp (including ~120 bp of adapter) was performed using a 4% Metaphor Agarose gel by Lonza (Cat: 50180). All libraries were pooled equimolar and sequenced 51 bp plus one 7 bp Index read on a Hiseq 2500 using Illumina v3 sequencing by synthesis (SBS) chemistry.

### Computational analysis of RNA-sequencing data

The quality control of RNA-sequencing data set was performed using the FASTQC tool [http://www.bioinformatics.babraham.ac.uk/projects/fastqc/]. The mapper utility of miRDeep2 v2.0.0.7^49^ was used to remove 3’ adapter sequences (AAGATCGGAAGAGCACACGTCT) of Illumina reads. Trimmed reads were then aligned using mapper utility to Bos taurus (UMD3.1 assembly, Ensembl release 84) reference genome with default parameters, except that the reads with < 17 nucleotides were discarded. miRBase (release 21)^50^ and miRDeep2.pl script were used to quantify expression levels of known microRNA species and to discover novel microRNA transcripts. The output of microRNA counts from miRDeep2.pl script were converted to library size normalized Count Per Million (CPM) values by using the cpm function of edgeR^51^ Bioconductor package. miRNAs that were expressed ≥ 1 log_2_ CPM in all samples were used in further analysis.

### Counting miRNA copy number and confirming vesicle-incorporated miR-375

The number of purified LDCVs was determined using Nanoparticle Tracking Analysis (NTA) (Malvern NanoSight LM10). In parallel, the concentration of miRNA extracted from LDCVs was determined using quantitative real-time PCR (qRT-PCR). A synthesized *Caenorhabditis elegans* microRNA (cel-miR-39) as the spike-in control was added to vesicle samples to normalize the RNA extraction efficiency. RNA was isolated using miRNeasy Mini Kit (Qiagen, Cat: 217004), then cDNA was made using miScript RT II (Qiagen, Cat:218161) according to the manufacturer’s protocol. A serial dilution of synthetic bta-miR-375 and cel-miR-39 was used to plot a standard curve to assess absolute miRNA quantification. The primers were UUUUGUUCGUUCGGCUCGCGUGA (Qiagen Cat: MS00053865) for bta-miR-375, and UCACCGGGUGUAAAUCAGCUUG (Qiagen Cat: MS00019789) for cel-miR-39.

For confirming vesicle-incorporated miR-375, 1 pmol of cel-miR-39 was mixed with 80 μg of purified LDCVs prior to RNase A treatment (10 μg/ml, 15 min, RT). RNase A selectively degrades vesicle-free miRNA whereas vesicle-incorporated miRNA is intact. 20 units of RNase inhibitor (ribonuclease inhibitor, Thermo Fisher Scientific, Cat: N8080119) was added before RNA extraction in order to inactivate RNase. As described above, RNA was isolated using miRNeasy Mini Kit, and then levels of cel-miR-39 and miR-375 with or without RNase A were determined by qRT-PCR followed by absolute miRNA quantification.

### Statistical analysis

All quantitative data are presented as means ± standard deviation (SD). Comparison between two groups were analyzed by two-tailed unpaired Student’s *t*-test. Statistical significance was set at *p* < 0.05.

## References

1. International Human Genome Sequencing, C. Finishing the euchromatic sequence of the human genome. Nature 431,931–45 (2004).

2. Frith, M.C., Pheasant, M. & Mattick, J.S. The amazing complexity of the human transcriptome. Eur J Hum Genet 13, 894–7 (2005).

3. Gomes, A.Q., Nolasco, S. & Soares, H. Non-coding RNAs: multi-tasking molecules in the cell. Int J Mot Sci 14, 16010–39 (2013).

4. Ha, M. & Kim, V.N. Regulation of microRNA biogenesis. Nat Rev Mot Cell Biot 15, 509–24 (2014).

5. Turturici, G., Tinnirello, R., Sconzo, G. & Geraci, F. Extracellular membrane vesicles as a mechanism of cell-to-cell communication: advantages and disadvantages. Am J Physiol Cell Physiol 306, C621–33 (2014).

6. Friedman, R.C., Farh, K.K., Burge, C.B. & Bartel, D.P. Most mammalian mRNAs are conserved targets of microRNAs. Genome Res 19, 92–105 (2009).

7. Iguchi, H., Kosaka, N. & Ochiya, T. Secretory microRNAs as a versatile communication tool. Commun Integr Biol 3, 478–81 (2010).

8. Turchinovich, A., Weiz, L. & Burwinkel, B. Extracellular miRNAs: the mystery of their origin and function. Trends Biochem Sci 37, 460–5 (2012).

9. Kosaka, N., Iguchi, H. & Ochiya, T. Circulating microRNA in body fluid: a new potential biomarker for cancer diagnosis and prognosis. Cancer Sci 101, 2087–92 (2010).

10. Iftikhar, H. & Carney, G.E. Evidence and potential in vivo functions for biofluid miRNAs: From expression profiling to functional testing: Potential roles of extracellular miRNAs as indicators of physiological change and as agents of intercellular information exchange. Bioessays 38, 367–78 (2016).

11. Chevillet, J.R. et al. Quantitative and stoichiometric analysis of the microRNA content of exosomes. Proc Nati Acad Sci USA 111, 14888–93 (2014).

12. Vickers, K.C., Palmisano, B.T., Shoucri, B.M., Shamburek, R.D. & Remaley, AT. MicroRNAs are transported in plasma and delivered to recipient cells by high-density lipoproteins. Nat Cell Bioi 13, 423–33 (2011).

13. Li, L. et al. Argonaute 2 complexes selectively protect the circulating microRNAs in cell-secreted microvesicles. PLoS One 7 e46957 (2012).

14. Xu, J., Chen, Q., Zen, K., Zhang, C. & Zhang, Q. Synaptosomes secrete and uptake functionally active microRNAs via exocytosis and endocytosis pathways. J Neurochem 124, 15–25 (2013).

15. Pegtel, D.M., Peferoen, L. & Amor, S. Extracellular vesicles as modulators of cell-to-cell communication in the healthy and diseased brain. Philos Trans R Soc Lond B Biol Sci 369 (2014).

16. Zhang, Q. et al. Selective secretion of microRNA in CNS system. Protein Cell 4, 243–7 (2013).

17. Park, C.K. et al. Extracellular microRNAs activate nociceptor neurons to elicit pain via TLR7 and TRPA1. Neuron 82, 47–54 (2014).

18. De Camilli, P. & Jahn, R. Pathways to regulated exocytosis in neurons. Annu Rev Physiol 52, 625–45 (1990).

19. Park, Y. & Kim, K.T. Short-term plasticity of small synaptic vesicle (SSV) and large dense-core vesicle (LDCV) exocytosis. Cell Signal 21, 1465–70 (2009).

20. Martin, T.F. The molecular machinery for fast and slow neurosecretion. CurrOpin Neurobiol 4, 625–32 (1994).

21. Winkler, H. The adrenal chromaffin granule: a model for large dense core vesicles of endocrine and nervous tissue. J Anat 183 ( Pt 2), 237–52 (1993).

22. Kasai, H., Takahashi, N. & Tokumaru, H. Distinct initial SNARE configurations underlying the diversity of exocytosis. Physiol Rev 92, 1915–64 (2012).

23. Park, Y. et al. Controlling synaptotagmin activity by electrostatic screening. Nat Struct MoI Biol 19, 991–7 (2012).

24. Park, Y. et al. Synaptotagmin-1 binds to PIP(2)-containing membrane but not to SNAREs at physiological ionic strength. Nat Struct Mol Biol 22, 815–23 (2015).

25. Simpson, R.J., Jensen, S.S. & Lim, J.W. Proteomic profiling of exosomes: current perspectives. Proteomics 8, 4083–99 (2008).

26. Alvarez-Erviti, L. et al. Lysosomal dysfunction increases exosome-mediated alpha-synuclein release and transmission. Neurobiol Dis 42, 360–7 (2011).

27. Stevanato, L., Thanabalasundaram, L., Vysokov, N. & Sinden, J.D. Investigation of Content, Stoichiometry and Transfer of miRNA from Human Neural Stem Cell Line Derived Exosomes. PLoS One 11, e0146353 (2016).

28. Fhaner, M.J., Galligan, J.J. & Swain, G.M. Increased catecholamine secretion from single adrenal chromaffin cells in DOCA-salt hypertension is associated with potassium channel dysfunction. ACS Chem Neurosci 4, 1404–13 (2013).

29. Winkler, H., Apps, D.K. & Fischer-Colbrie, R. The molecular function of adrenal chromaffin granules: established facts and unresolved topics. Neuroscience 18, 261–90 (1986).

30. Lawrie, C.H. et al. Detection of elevated levels of tumour-associated microRNAs in serum of patients with diffuse large B-cell lymphoma. Br J Haematol 141, 672–5 (2008).

31. Mitchell, P.S. et al. Circulating microRNAs as stable blood-based markers for cancer detection. Proc Natl Acad Sci U S A 105, 10513–8 (2008).

32. Rayner, K.J. & Hennessy, E.J. Extracellular communication via microRNA: lipid particles have a new message. J Lipid Res 54, 1174–81 (2013).

33. Thureson-Klein, A.K. & Klein, R.L. Exocytosis from neuronal large dense-cored vesicles. Int Rev Cytoil 121, 67–126 (1990).

34. Poy, M.N. et al. miR-375 maintains normal pancreatic alpha- and beta-cell mass. Proc Natl AcadSci USA 106, 5813–8 (2009).

35. Bhinge, A. et al. MiR-375 is Essential for Human Spinal Motor Neuron Development and May Be Involved in Motor Neuron Degeneration. Stem Cells 34, 124–34 (2016).

36. Abdelmohsen, K. et al. miR-375 inhibits differentiation of neurites by lowering HuD levels. Mot Cell Biol30, 4197–210 (2010).

37. Erener, S., Mojibian, M., Fox, J.K., Denroche, H.C. & Kieffer, T.J. Circulating miR-375 as a biomarker of beta-cell death and diabetes in mice. Endocrinology 154, 603–8 (2013).

38. Yin, J., Hou, P., Wu, Z., Wang, T. & Nie, Y. Circulating miR-375 and miR-199a-3p as potential biomarkers for the diagnosis of hepatocellular carcinoma. Tumour Biol36, 4501–7 (2015).

39. Denk, J. et al. MicroRNA Profiling of CSF Reveals Potential Biomarkers to Detect Alzheimer’s Disease. PLoS One 10, e0126423 (2015).

40. Sanek, N.A. & Young, W.S. Investigating the in vivo expression patterns of miR-7 microRNA family members in the adult mouse brain. Microrna 1, 11–8 (2012).

41. Malmevik, J. et al. Identification of the miRNA targetome in hippocampal neurons using RIP-seq. Sci Rep 5, 12609 (2015).

42. de Chevigny, A. et al. miR-7a regulation of Pax6 controls spatial origin of forebrain dopaminergic neurons. Nat Neurosci 15, 1120–6 (2012).

43. Le, M.T. et al. MicroRNA-125b promotes neuronal differentiation in human cells by repressing multiple targets. Moi Cell Biol 29, 5290–305 (2009).

44. Sawada, K. et al. Identification of a vesicular nucleotide transporter. Proc Natl Acad Sci U S A 105, 5683–6 (2008).

45. Pobbati, A.V., Stein, A. & Fasshauer, D. N- to C-terminal SNARE complex assembly promotes rapid membrane fusion. Science 313, 673–6 (2006).

46. Park, Y. et al. alpha-SNAP interferes with the zippering of the SNARE protein membrane fusion machinery. J Biol Chem 289, 16326–35 (2014).

47. Smith, A.D. & Winkler, H. A simple method for the isolation of adrenal chromaffin granules on a large scale. Biochem J 103, 480–2 (1967).

48. Ahmed, S., Holt, M., Riedel, D. & Jahn, R. Small-scale isolation of synaptic vesicles from mammalian brain. Nat Protoc 8, 998–1009 (2013).

49. Friedlander, M.R., Mackowiak, S.D., Li, N., Chen, W. & Rajewsky, N. miRDeep2 accurately identifies known and hundreds of novel microRNA genes in seven animal clades. Nucleic Acids Res 40, 37–52 (2012).

50. Griffiths-Jones, S., Grocock, R.J., van Dongen, S., Bateman, A. & Enright, A.J. miRBase: microRNA sequences, targets and gene nomenclature. Nucleic Acids Res 34, D140–4 (2006).

51. Robinson, M.D., McCarthy, D.J. & Smyth, G.K. edgeR: a Bioconductor package for differential expression analysis of digital gene expression data. Bioinformatics 26, 139–40 (2010).

